# Convergent enrichment of Gammaproteobacteria along *Aedes aegypti* development across different breeding sites

**DOI:** 10.1101/2025.09.13.676009

**Authors:** Aboubakar Sanon, Lander De Coninck, Lanjiao Wang, Athanase Badolo, Jelle Matthijnssens, Katrien Trappeniers, Leen Delang

## Abstract

**Background:** *Aedes aegypti* mosquitoes are the main vector of pathogens like dengue virus and chikungunya virus. The immature life stages of mosquitoes share the same habitat with a variety of microorganisms in aquatic environments. To better understand the microbial diversity in field-derived *Ae. aegypti*, we analysed simultaneously collected larvae, pupae, and freshly emerged adults from Burkina Faso together with their breeding water via 16S rRNA gene sequencing.

**Results:** We observed a decrease in bacterial diversity and load across the mosquito life stages. At the phylum level, a strong increase in relative abundance of Proteobacteria was found along the mosquito stages. The same 40 amplicon sequence variants were consistently found as most abundant in the adults, regardless of the sample collection site, and all belonged to the Gammaproteobacteria. Our data suggest that these bacteria were not randomly derived by chance from the environment in the mosquito but rather deposited by a female mosquito during oviposition, a transmission route recently coined as “diagonal transmission”. Indeed, our results indicated that there is a selection of Gammaproteobacteria from the breeding water and that these bacterial members are further maintained from larvae to adults.

**Conclusion:** This study provided new data on the microbiome composition of field-collected *Ae. aegypti*, contributing to an enhanced understanding of the origin and colonization route of the mosquito microbiome, potentially via a diagonal transmission route.

## Introduction

*Aedes aegypti* mosquitoes are the main vector of pathogenic arboviruses like dengue virus, chikungunya virus, and Zika virus in tropical and subtropical areas [1]. *Ae. aegypti* thrives mostly in urban environments, where the vector finds suitable conditions to survive and adapt to growing urbanization [2, 3]. Humans are the most important host for this mosquito species [2]. People in tropical and subtropical regions are the most affected by arbovirus infections compared to other, more temperate regions, although higher latitude areas are not exempt from these infections because of global travel and climate warming [4]. Currently used vector control methods are often inefficient, exemplified by *Ae. aegypti* exhibiting a high level of resistance to insecticides [5]. As a result, *Ae. aegypti* is still expanding to new environments due to growing urbanization, deforestation, and global warming [6]. New strategies targeting the vector are being explored to fight arbovirus-induced diseases.

The insect microbiota plays an important role in health and fitness, for instance by providing nutritional and defence benefits for the insect host [7–11]. Particularly during mosquito development, resident bacterial organisms have an important nutritional role [12]. Furthermore, the microbiota also affects vector competence, *i.e.* the intrinsic ability of an arthropod vector to transmit an infectious agent [13–15]. This mosquito microbiota has become an interesting target to reduce both mosquito populations as well as vectorial capacity for important mosquito-borne diseases [16, 17]. In order to further exploit the mosquito microbiota as a new tool for vector control in the field, a better and more complete understanding of the microbial composition and colonisation process in mosquitoes is needed, particularly for field-caught mosquitoes.

*Aedes* mosquitoes are holometabolous insects, meaning they proceed to a complete metamorphosis and undergo a developmental process with four distinct stages: egg, larva, pupa, and adult. The egg hatches in aquatic habits, followed by subsequent developmental stages of different instar larval stages and a pupal stage, before emerging as adult. Immature mosquitoes (larvae and pupae) and adult mosquitoes are confined to aquatic and terrestrial habitats respectively. After mating, the female will bite a vertebrate host to acquire a blood meal containing essential amino acids needed for egg maturation [8]. Each female mosquito lays multiple egg clutches during its life cycle and each clutch ranges from ten to hundreds of eggs [18]. A crucial step in the mosquito life cycle is the deposition of eggs by the gravid female as it determines the fate of the progeny and thus oviposition behaviour is thought to be under strong selection pressure. Mosquito species are known to have a preference for specific breeding habits, depending on certain characteristics such as sunlight exposure, salinity of the water, stream flow, organic matter, predators, etc. [19]. *Ae. aegypti*, for instance, will oviposit in both natural and artificial containers, while some *Culex* species prefer to lay eggs in freshwater pools with high organic matter [19]. Urban areas are favourable habitats for *Aedes* mosquitoes due to the availability of diverse sugar sources and abundant blood meals, less predators, and multiple water-holding containers needed for egg deposition and larval development [20, 21]. Biotic and abiotic factors in the water can drive the selection of breeding sites by the mosquito female [22–25]. The aquatic environments of larvae and pupae are often rich in organic matter acquired from the soil, vegetation, animal cadavers, etc. and suited to promote the growth of a diversity of microorganisms [26]. Larvae obtain organic compounds from their aquatic habit using different feeding strategies including filtering, suspension feeding, or predation [26]. Pupae on the other hand, are a non-feeding stage of the mosquito life cycle and do not ingest any food from their environment.

The bacterial communities of *Ae. aegypti* span multiple phyla, with Proteobacteria typically dominating alongside Actinobacteria, Firmicutes, and Bacteroidetes in both aquatic and adult stages [14, 27]. Within these communities, adult midguts and crops frequently harbour members of the families Acetobacteraceae and Weeksellaceae, with genera such as *Asaia*, *Elizabethkingia*, and *Tanticharoenia* often being highly abundant [28]. Broader surveys and reviews further show that gut and tissue microbiota of *Aedes* spp. commonly include *Enterobacter*, *Klebsiella*, *Pantoea*, *Serratia*, *Pseudomonas*, *Sphingomonas*, *Cupriavidus*, *Escherichia–Shigella*, *Enterococcus*, and *Bacillus*, although relative abundances vary with geography, rearing conditions, and life stage [25, 29].

In the past years, more studies investigated the role of microorganisms colonizing the mosquito habitat. Interestingly, the choice of oviposition site by *Ae. aegypti* was shown to be influenced by the microbial communities in the water [30–32]. *Aedes* females find a suitable site to lay their eggs, at least partially, using microbial emitted chemicals [33]. Although other studies have described a transstadial transmitted gut microbiome in *Anopheles* mosquitoes [34–36], the exact origin of how microbial communities colonize mosquitoes and which species are maintained across different life stages remains not completely understood. Mosquitoes, like other insects, can acquire their microbiota via vertical transmission, *i.e.* direct transmission from parent to offspring, or horizontal transmission, *i.e.* transfer of bacteria between individuals that are not in a parent-offspring relationship [37]. Horizontal transmission often occurs through contact, feeding, mating, or shared environments. Larvae, for instance, acquire their resident microbes by feeding on environmental aquatic organic detritus, unicellular filamentous algae, bacteria, and other microorganisms [14, 38, 39]. Studies on the microbiome composition in different life stages of *Ae. aegypti* and their breeding sites can therefore provide insights on the origin of the mosquito microbiota and its influence on *Ae. aegypti* fitness and development. Only a limited number of studies have reported on transstadial microbiome analyses for *Ae. aegypti,* including one report studying lab mosquitoes. Most reports on field mosquitoes do not include all mosquito life stages (for instance only water and larvae) [27, 40–43].

In this study, we described (i) the variation of microbial communities of field-collected *Ae. aegypti* mosquitoes from Burkina Faso at different developmental stages (larvae, pupae, adult) and of the breeding water and (ii) determined whether breeding container (in which the mosquitoes breed; plastic or tire) or sampling sites (geographical location; urban or peri-urban) can influence bacterial communities of water and mosquito samples.

## Methodology

### Mosquito collection

In total, 48 individual samples (12 *Ae. aegypti* individuals across three life stages and 12 water samples) were collected in Ouagadougou, Burkina Faso during the rainy season of 2022 (**Figure 1A**). In brief, samples were collected from two sampling sites (urban and peri-urban) and from two types of breeding material (plastic containers and tires) in each sampling site (**Figure 1B and C**). The two sample sites were chosen based on their urbanization type. 1200 Logements is an urban area where the roads are paved and there is a modern system for wastewater evacuation. Toudbweogo lies in a transitional zone between the modern urban centre of Ouagadougou and surrounding rural villages. The area lacks a wastewater evacuation system, has limited utilities, and unpaved roads; it is therefore classified as peri-urban. For each breeding material, mosquito samples from different life stages were collected (third or fourth instar larvae, pupae, and freshly emerged adults) as well as a sample of the water present in the breeding material (i.e. three biological replicates per condition). The two sampling sites were located approximately 10 km from each other. Water, larvae, and pupae were taken with a sterile Pasteur pipette from the original water of the breeding site and transferred into sterile Eppendorf tubes. At the same time or the next day, newly emerging adults from collected pupae were aspirated in the lab’s insectary with a mouth aspirator, killed by a cold shock, and stored at -80°C. All samples were shipped on dry ice to Belgium for DNA extraction and 16S rRNA gene sequencing.

**Figure 1:**
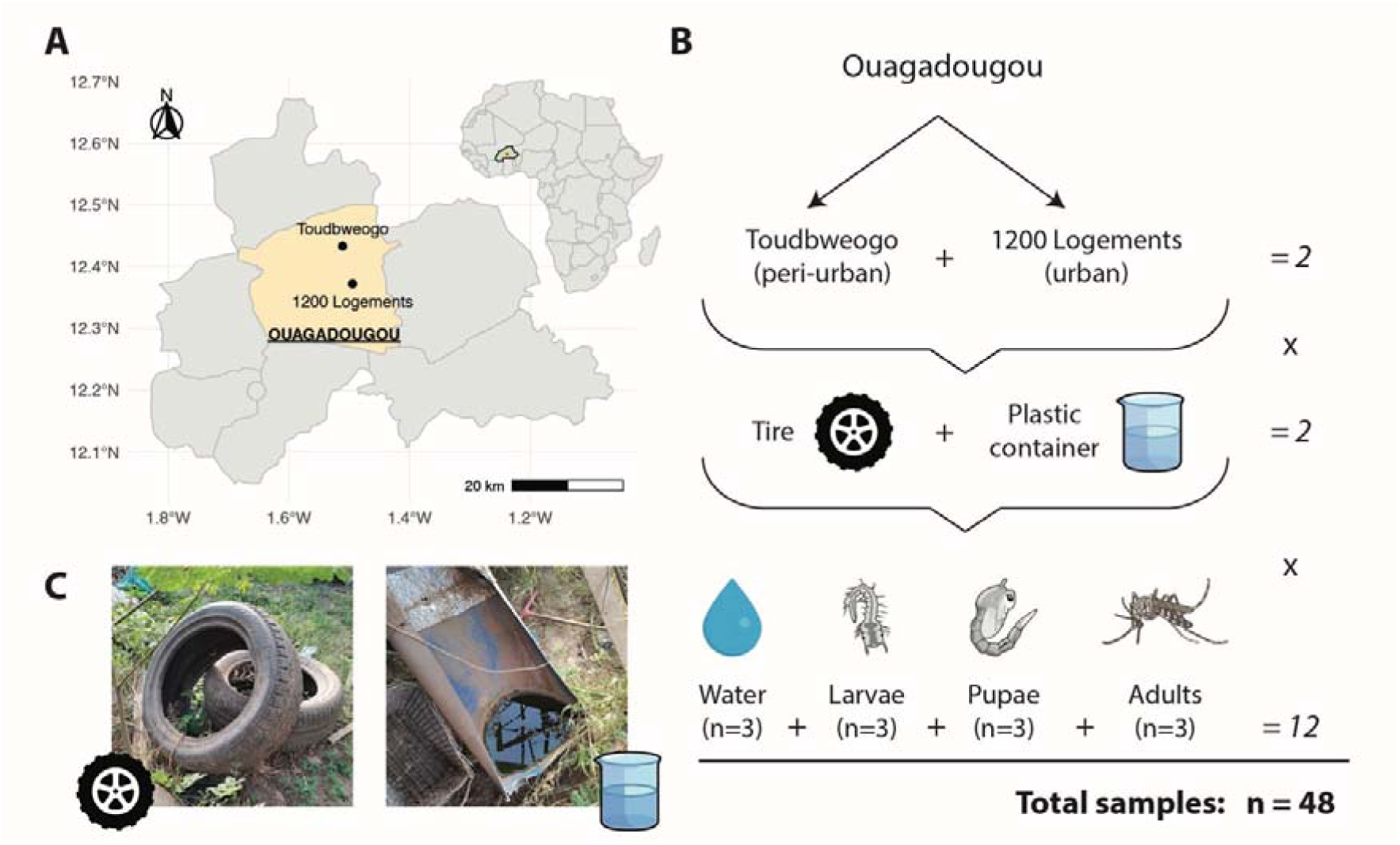
**(A)** Map of the Kadiogo region showing the Ouagadougou district in yellow. The two collection sites are indicated with black dots. The inset map shows the African continent with Burkina Faso coloured green and the Kadiogo region red. **(B)** Overview of the sampling collection. Samples were collected from two sampling sites (urban and peri-urban) and from two types of breeding material (plastic containers and tires) in each sampling site. For each breeding material, mosquito samples from different life stages were collected (larvae, pupae, and freshly emerged adults) as well as a sample of the water present in the breeding material, all in triplicate. A total of 48 samples were collected. **(C)** Representative images of the tire and plastic container breeding materials.

### DNA extraction

DNA was extracted from the mosquito samples under a laminar flow in sterile conditions. Larvae (third or fourth instars), pupae, and adults were rinsed in PBS for 3 minutes, then in 70% ethanol, and finally in sterile water to sterilize their surface and remove any bacteria coming from the external environment. Individual samples, whole larvae, pupae, and female adults (whole bodies) were suspended in tubes with ceramic beads (Precellys) containing 150 µL of PBS, homogenized for 1 min, and centrifuged (12000 rpm for 4 min). The homogenate was treated with lysozyme and proteinase K and incubated at 56°C for 5 min to completely lyse the tissue. Then, the homogenate was transferred in an Eppendorf tube and DNA was extracted using the DNeasy 96 Blood & Tissue DNA extraction kit (Qiagen). An aliquot of the water samples was taken from the original breeding site after pipetting up and down to mix the water sample and 500 µL of this debris was put into Eppendorf tubes and centrifuged at 5000 g for 30 minutes to pellet microbes in the water. The supernatant was discarded, and the resuspended pellet was used for DNA isolation following the same protocol for mosquito DNA extraction as described above. The final DNA was eluted in 100 µL of sterile elution buffer (100 mM Tris-HCl) and quantified using Nanodrop. The samples were shipped on dry ice to Macrogen Netherlands for 16S rRNA gene sequencing. The V3-V4 hypervariable region of the bacterial 16S rRNA gene was PCR amplified, and the sequencing was carried out on the Illumina MiSeq system with a 300bp paired-end run. Furthermore, 6 negative controls (PBS) which were processed together with the 36 mosquito and 12 water samples following the same DNA extraction protocol were sequenced as blank controls.

### 16S rRNA gene sequencing data processing and s

We used the DADA2 pipeline v1.16 with default parameters specified by the authors to identify amplicon sequence variants (ASVs) [44], which are superior to operational taxonomic units (OTUs) in terms of taxonomic resolution. Each ASV was subsequently assigned a taxonomic identity using DECIPHER’s IDTAXA [45] with the SILVA SSU r138 training data set, and generated count data for each ASV. The code used for sequence processing, bioinformatics, statistical analyses, and the data can be found at https://github.com/absanon/2024_XXX_MICROBIOME_LIFE_STAGE. We imported the ASV data from DADA2 into a phyloseq object for bacterial community analyses and visualization in R. As an additional quality control step, we used the decontam R package [46] with the “prevalence” method and a stringent probability threshold of 0.5 to identify and remove ASVs as contamination. Alpha diversity metrics like the Shannon diversity index and the Simpson index were assessed with the “vegan” package [47] by performing a rarefaction analysis [48] on the original abundance table (1000 iterations sampled to 47,006 reads, which was the minimum sequencing depth). We used the Wilcoxon test with the Benjamini-Hochberg FDR correction for multiple comparisons to test for differences in the mean alpha diversity between sample types (water *vs* larvae *vs* pupae *vs* adults). Bray-Curtis dissimilarities between samples were calculated using vegan’s avgdist function, again sampling the original abundance table to 47,006 reads per sample for 1000 iterations. A principal coordinate analysis (PCoA) was performed on the Bray-Curtis dissimilarities with the “ape” package [49]. To identify the impact of the different metadata variables (mosquito life stage, location and breeding material) on the bacterial community composition of the samples, we used a distance-based redundancy analysis (dbRDA). Using an adapted version of https://github.com/raeslab/RLdbRDA allowing for a custom distance matrix as input, this procedure consisted of two steps. First, the effect of each explanatory variable on community composition was evaluated individually using the capscale function implemented in the “vegan” package [47] (univariate analysis). Subsequently, variables identified as significant in this initial assessment were subjected to a stepwise selection procedure to handle correlated metadata variables and to determine which of them exert the strongest influence on community composition (multivariate analysis).

### Analysis of public 16S data from field *Aedes aegypti*

To confirm the alpha diversity and bacterial composition results, we searched through the literature for other studies that did 16S sequencing on field-collected *Ae. aegypti* samples with at least two different life stages. We identified four other studies that had a somewhat similar methodology compared to our study (see **Table 1**) and after downloading the public data, we performed the same data analysis as on our samples to enable comparisons between the datasets (see above).

**Table 1:** Overview of studies with public 16S data on field Ae. aegypti samples across multiple life stages.

| Study | Journal | 16S regions | Country | Life stages | BioProject |
| --- | --- | --- | --- | --- | --- |
| Bennett et al. (2019) | Scientific reports | V4 | Panama | Larvae, adults | PRJNA523634 |
| Hery et al. (2021) | Microbial Ecology | V6-V8 | Guadeloupe/French Guiana | Water, larvae | PRJNA600474 |
| Rodpai et al. (2023) | Insects | V3-V4 | Thailand | Water, larvae, adults | PRJNA919511 |
| Hernández et al. (2024) | PLoS One | V3-V4 | Mexico | Eggs, larvae, adults | PRJNA836318 |

### Absolute quantification of 16S copies with qPCR

A qPCR was performed to obtain the absolute number of 16S copies in each sample. Total bacterial DNA load was quantified using quantitative PCR (qPCR) by amplification of the bacterial 16S rRNA gene using universal 16S primers targeting the V3-V4 region. A total reaction volume of 20 µL, containing 5 µL sample and 15 µL master mix, was used. Per reaction, the master mix consisted of 10 µL SYBR green reaction mix (ITAQ UNIV SYBR, Bio-Rad), 0.4 µL 10 µM forward primer (IDT), 0.4 µL 10 µM reverse primer (IDT), and 4.2 µL nuclease-free water. Following 16S primer sequences were used: forward primer 5’-TCCTACGGGAGGCAGCAGT-‘3 and reverse primer 5’- GGACTACCAGGGTATCTAATCCTGTT-‘3 [50]. Reactions were performed in the QuantStudio 5 Real-Time PCR System (Applied Biosystems, Thermo Fisher Scientific). The amplification conditions were 1 cycle of 10 min at 95°C, 40 cycles of 15 sec at 95°C, and 40 cycles of 15 sec at 55°C. To quantify the bacterial genome copies in each sample, a 1:10 standard curve was made using a 16S DNA gene block (IDT).

Absolute ASV abundances per sample were obtained by multiplying their relative abundance, derived from the NGS data, by the total number of 16S copies/µl from the qPCR results.

## Results

### Bacterial richness and composition are determined by mosquito life stage

In total, 48 samples were collected in the capital of Burkina Faso consisting of 12 water samples and 36 mosquito samples. Different mosquito life stages were collected (larvae/pupae/newly emerged adult mosquitoes) for two sampling sites (urban/peri-urban), and two types of breeding material (tire/plastic container), all in triplicate. The DNA of the mosquito samples (larvae, pupae, and whole body for the adults) as well as the water samples was extracted and analysed using 16S rRNA gene sequencing. On average, 114,143 sequencing reads were generated for each sample with a minimum of 84,548 and a maximum of 147,901 reads. After removing chimeras and filtering out contamination, a total of 24,141 ASVs were identified across the 48 samples with the DADA2 pipeline.

To evaluate the bacterial community diversity, the alpha diversity was calculated based on the obtained ASVs and compared between the mosquito and water samples. A decreasing bacterial alpha diversity was observed from the water samples over the larvae and pupae towards the adult mosquito samples. Although the alpha diversity between the water and larval stages was not significantly different, a statistically significant decrease was found in the pupal and adult stages (P<0.05; **Figure 2A**). We did not observe any difference in alpha diversity between both breeding materials and locations (see **Supplemental Figure 1**) Overall, the bacterial community diversity decreased according to the mosquito developmental stages. These results were mostly confirmed by the public data analysis (see **Supplemental Figure 2**), only in the Rodpai *et al.* dataset [43] there was no clear decrease in alpha diversity across the different life stages likely due to the low number of replicates. On the other hand, all other datasets showed a decrease of alpha diversity across the available life stages and these were significant for both the Hery and Hernández studies [27, 42] (see **Supplemental Figure 2**).

**Figure 2:**
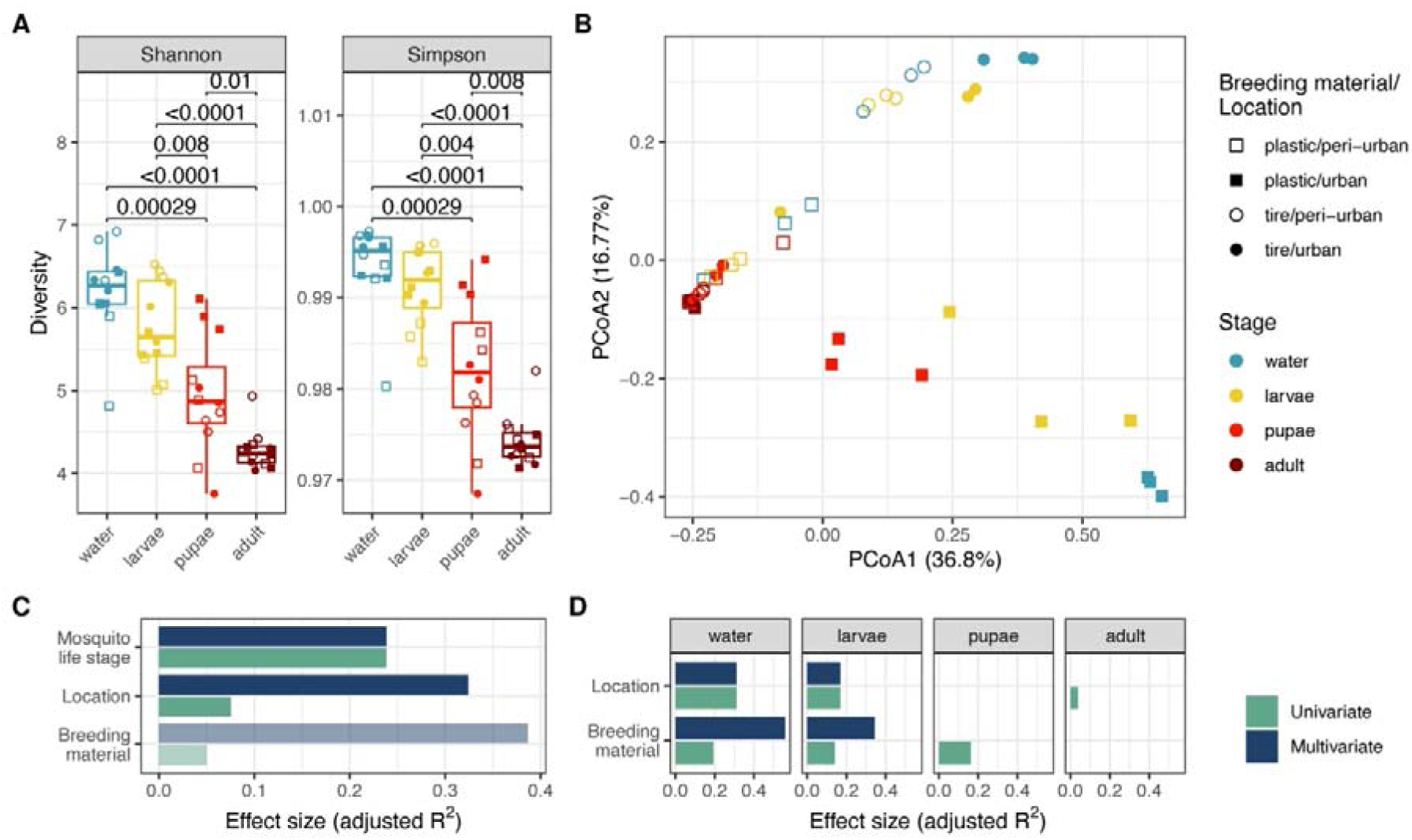
**(A)** Alpha diversity measures (Shannon and Simpson) between the water samples and the different mosquito developmental stages. The number on top of the boxplots represents the respective adjusted p-value of the comparisons between life stages. Wilcoxon tests with Benjamini-Hochberg FDR correction were used to compare the diversity of the bacteria between the water community and the different mosquito developmental stages. **(B)** PCoA plot showing the clustering in bacterial communities of mosquito developmental stages, sampling site, and breeding material. **(C)** The effect size of developmental stage, location and breeding material on the bacterial composition calculated using dbRDA. Variables are ordered according to effect size of significant covariates selected by redundancy analysis (navy bars) compared with individual effect sizes assuming independence (green bars). Light coloured bars indicate a non-significant multivariate effect of that variable. **(D)** Effect size of location and breeding material on the variance in the developmental stage of mosquito microbiota composition. The variables were given a fixed order in this plot with location preceding breeding material. No bars indicate a non-significant effect of the variable on an independent test, making the multivariate test unnecessary.

To study the impact of the mosquito life stages (larvae, pupae, adults), the sampling location (urban, peri-urban), and the type of breeding material (tire, plastic) on the microbiome composition, a principal coordinate analysis (PCoA) was performed. Again, the PCoA showed that the bacterial communities of the water samples as well as the early mosquito life stages were highly diverse, whereas the microbial composition of the freshly emerged adult mosquitoes was very similar and irrespective of the sampling site or breeding material (**Figure 2B**). In addition, the water and immature stages of plastic containers from the urban site displayed a large variation among the samples (**Figure 2B**). The impact of the life stages, the type of breeding material, and the sampling site on the microbial communities was investigated using a distance-based redundancy statistical analysis showing the influence of these three factors on the microbiome composition. Mosquito life stage explained the majority (24%) of the microbial variation while sampling location and the type of breeding material explained respectively 8% and 5% of the microbial variation (**Figure 2C**). In the multivariate analysis, only the mosquito life stage and the location were explanatory with respect to the microbiome composition, without additional information from the breeding material. Next, we further analysed the effect of the sampling site and the type of breeding material on the microbial variation within each mosquito life stages. The redundancy analysis identified that in the early life stages, location and breeding material explained a large amount of the bacterial variation with respectively 18% and 15% in larvae. While the breeding material still explained 18% of the variation in pupae, neither location nor the breeding material showed a major contributing role on the bacterial composition in adults (**Figure 2D**).

### Gammaproteobacteria dominate the *Aedes aegypti* microbiome

Next, the bacterial composition was investigated at the family level for the mosquito life stages, types of breeding material, and sampling sites (**Figure 3**). For this, only bacterial phyla with a relative abundance of >1% are shown, otherwise they were grouped in ‘Others’ together with unclassified ASVs. In addition, only bacterial families with an abundance higher than 5% in at least one sample, or at least the most abundant family per phylum, are shown. Remaining bacterial families were pooled together based on their phylum. In total, we found 9 phyla with a relative abundance higher than 1%, which included 33 bacterial families (**Figure 3**). The water samples showed a large variability of different bacterial phyla with the Proteobacteria, Actinobacteriota, Bacteriodota, and Firmicutes as most abundant phyla (**Figure 3**). Across the life stages, the abundance of the Actinobacteriota, Bacteriodota, and Firmicutes decreased gradually along the developmental axis. Eventually, the Proteobacteria, more specifically the Gammaproteobacteria class, became dominant in all adult mosquitoes, occupying 80% to 90% of the relative abundance in each adult sample, regardless of their sampling location or breeding material (**Figure 3** and **Supplemental Figure 3**).

**Figure 3:**
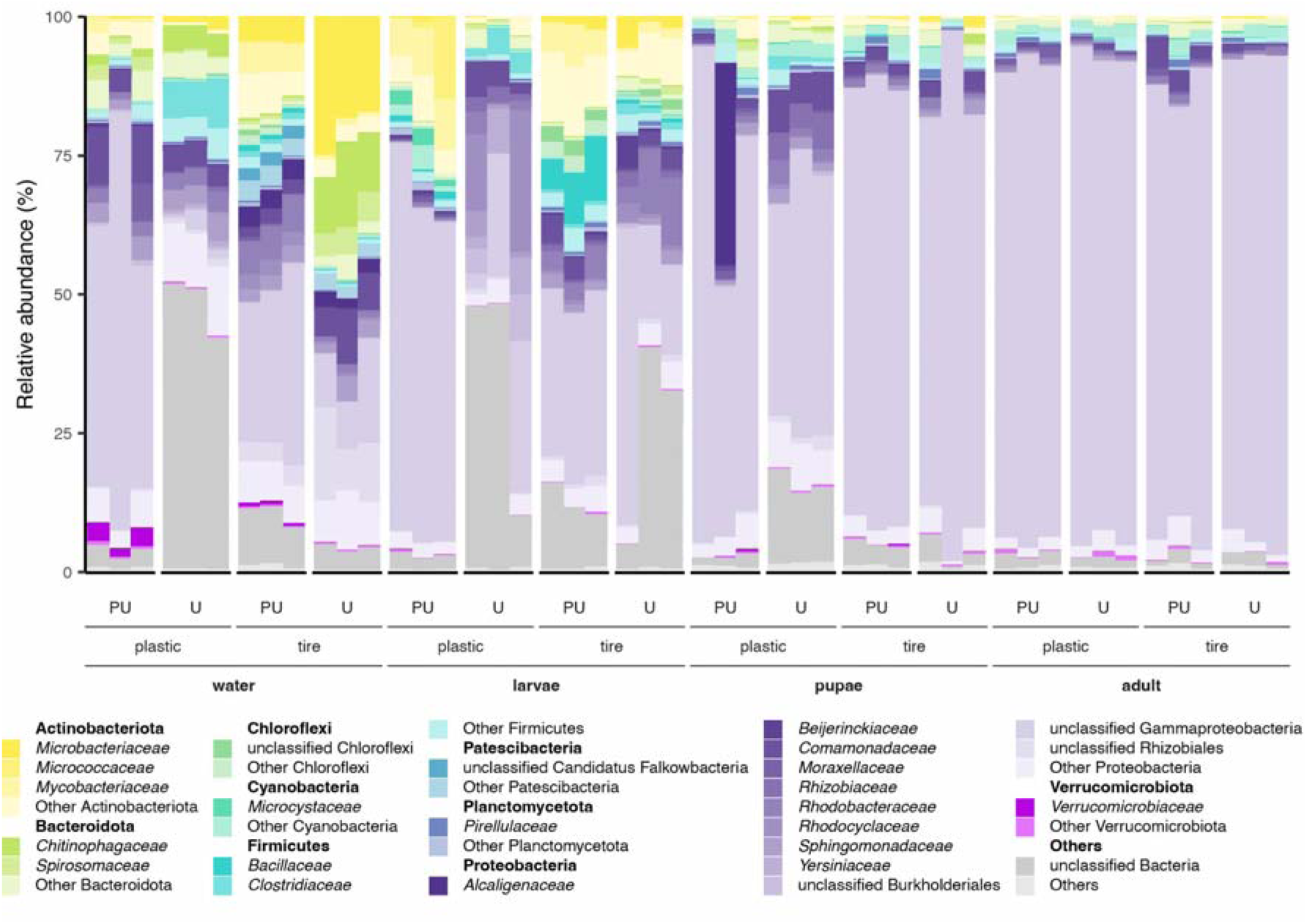
Relative abundance of identified bacterial families shown with different shades of the same colour per phylum. Only phyla with a relative abundance >1% are shown, and within each phylum only families with a relative abundance >5% in at least one sample are shown individually. Unclassified bacteria and phyla with a relative abundance <1% are shown as ‘Others’. Samples are grouped by biological replicates. U: urban, PU: peri-urban.

### Convergence to low divergent gut microbial composition in adult mosquitoes, irrespective of sampling site and breeding material

Interestingly, from the wide diversity of ASVs in the water, consistently the same set of 40 ASVs were found to be most abundant in the adult mosquitoes, regardless of the sampling site or breeding material (**Figure 4A** and **Supplemental Figure 4**). To avoid overplotting, we zoomed-in on the top 15 most abundant bacterial ASVs per mosquito life stage, sampling site, and breeding material (**Figure 4B and 4C**). In contrast to larvae, hence the major route of bacterial introduction in mosquitoes is halted during this developmental stage. Our data suggest that a small set of bacterial ASVs is further selected inside the mosquito during the later stages of the mosquito development. As a consequence, many bacterial ASVs abundant in the water and larval stages completely disappeared in the pupae and adult stages, explaining the decrease in microbial diversity with the subsequent life stage, and secondly the increase in relative abundance of the same set of 15 ASVs in later life stages, which represent approximately 60% of the bacterial load in the adult samples (**Figure 4C**).

**Figure 4:**
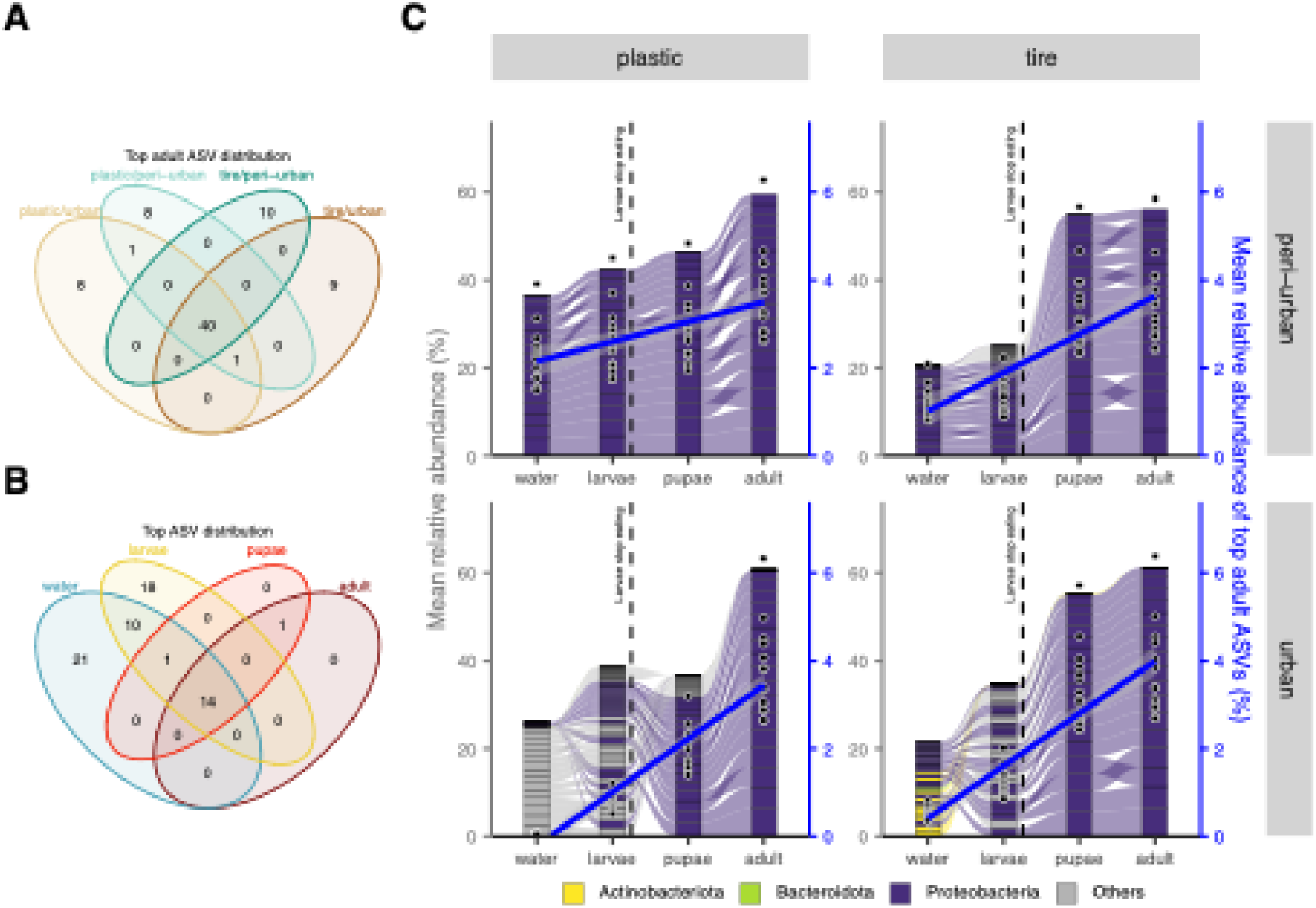
**(A)** Venn diagram displaying the distribution of the top 50 ASVs in the adults across the different locations and breeding materials showing that adult mosquitoes share the same most abundant 40 ASVs regardless of their collection site. **(B)** Venn diagram showing the sharing of the top 15 ASVs across developmental stages. ASVs were selected separately per location and breeding material, resulting in >15 ASVs appearing in the earlier developmental stages. **(C)** Alluvial plot showing the changes of the mean relative abundance of the top 15 ASVs across developmental stage (x-axis), location (y-axis split) and breeding material (x-axis split). For each location, breeding material and developmental stage the 15 most abundant ASVs were selected, and all are shown in each panel of the plot. The second y-axis (blue) indicates the mean relative abundance of the 15 top ASVs in the adult mosquitoes with a linear regression (blue line and black dots) showing the increase of their relative abundance across mosquito developmental stages.

The increase in relative abundance of these ASVs can be explained by either an overgrowth of these specific bacteria (most likely resulting in a stable or increasing bacterial load) or otherwise by a die-off of the other bacterial ASVs (most likely resulting in a decreasing bacterial load), with both resulting in an increase of the relative abundance of the remaining ASVs. To further investigate this, we performed a 16S qPCR analysis in the same DNA samples that were used for the 16S rRNA gene sequencing, allowing to evaluate the bacterial load and combine this with the relative 16S data. On average, water, larvae, pupae, and adult samples contained 35103, 6854, 524, and 39 16S copies/µl, respectively. The bacterial load thus decreased along the different developmental stages with the adult mosquitoes having the lowest total number of bacteria (**Figure 5A** and **Supplemental Figure 5**), corroborating with the decreasing alpha diversity pattern shown in **Figure 2A**. In fact, the absolute copy number of the 15 most abundant ASVs, obtained by multiplying the relative abundance from the NGS data with the absolute copy number from the 16S qPCR, in the adults is even higher in the water and larval stages (**Figure 5B**), suggesting that the developing mosquito body is a challenging environment for bacteria to thrive and only a few bacteria can survive here.

**Figure 5:**
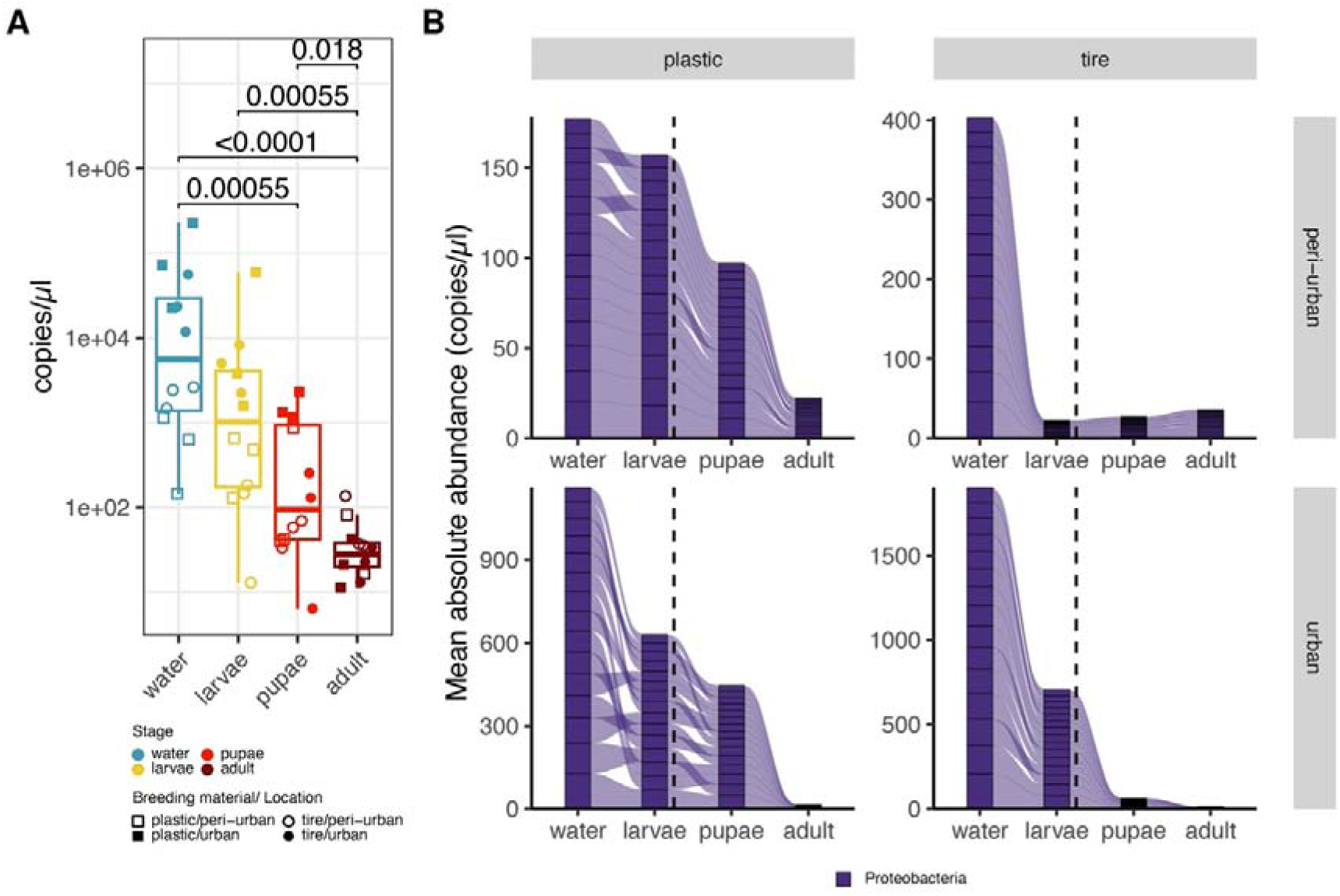
**(A)** Box plots showing the 16S copies/µl per sample derived from a SYBR qPCR with universal 16S primers. The number on top of the boxplots represents the respective adjusted p-value of the comparisons between life stages from Wilcoxon tests with a Benjamini-Hochberg FDR correction. **(B)** Alluvial plot showing the average absolute abundance of the 15 most abundant ASVs in the adult mosquitoes across the developmental stages, location and breeding material. The absolute abundance for each ASV per sample was calculated by multiplying the per-sample 16S copies/µl from the qPCR results with the relative abundance of these ASVs calculated from the NGS data. The dotted line specifies the transition from larvae to pupae indicating the time when environmental bacteria can no longer enter the larvae through filter feeding.

## Discussion

This study described the bacterial communities of *Ae. aegypti* from the capital city of Burkina Faso according to mosquito life stages, breeding material, and sampling sites from urban and peri-urban locations. In general, we observed a high diversity and bacterial load in water samples compared to the different mosquito developmental stages. Bacterial diversity was similar between the water and the larvae, reflecting the abundant inflow of environmental bacteria during the filter feeding activity of the larvae. However, once the pupae stage is reached, this filter feedings stop and a strong selection towards a small number of bacterial ASVs was observed in the adult stage. This observation can be explained by the process of metamorphosis. As larvae shed their epithelial gut lining with each moulting and at pupation excrete their gut contents, this will lead to a decrease in microbial abundance as shown by Moll *et al.* [51].

Our data showed significantly distinct bacterial compositions between pupae and adults, both with respect to diversity (**Figure 2A**) and bacterial load (**Figure 5A**), which is in line with previous observations [22, 25], but in contrast to the study of Bennet and colleagues [40], which described few differences between bacterial communities of larvae and adult *Aedes* species despite distinct habitat features like geographical location, type of breeding material, and associated environmental variables of the water. We also showed that the sampling site (urban or peri-urban) or the breeding material (plastic containers or tires) had an impact on the bacterial composition of the water and the larvae, but not on the pupae and the adult mosquitoes (**Figure 2D**). Of interest was the observation that the water, larvae and pupae samples collected in the urban area in the plastic containers showed a greater number of (unclassified) ASVs compared to other collection sites and the adult stages. This might be explained by a greater contamination of this environment due to human activities.

As shown in **Figure 4C**, the most abundant ASVs observed in adult mosquitoes were already present in the breeding water. This suggests that mosquitoes acquire their microbiota from their breeding water, which is in line with previous findings [23, 25, 52]. The question remains whether the breeding site shapes the bacterial community in the mosquito or, the other way around, whether female mosquitoes inoculate the breeding sites with relevant bacteria (eg. by excreting them from the gut) before, during or after deposition of their eggs in the aquatic environment as shown by Mosquera *et al.* [53]. The fact that across the different locations and the different breeding materials we consistently find the same most abundant ASVs in the adults, strongly suggests the latter. Particularly as ASV classification is very specific (a single nucleotide difference makes it a different ASV), the chance of randomly finding the same set of bacteria from so many different environmental sources should be extremely rare. This means that these bacteria were rather derived from a mosquito female and deposited in the breeding water from which the larvae acquired them via filter feeding. This mode of transmission does not directly fit into the two mainly known categories of microbial transmission (ie. vertical and horizontal transmission) in mosquitoes. Recently, the concept of ‘diagonal transmission’ was proposed for viral transmission by Hamel *et al*. [54]. They showed in their study the ability of West Nile virus to infect mosquito larvae through the adult excreta. We hypothesize that a similar route could apply for the microbiome as well, where microorganisms of one parent are deposited to the environment and transmitted to the offspring of the same or another parent (see **Supplemental Figure 10**). Diagonal transmission differs from vertical transmission as the microbiome of a parent can go to a different offspring, while vertical transmission comprises a direct line from parent to its offspring. Diagonal transmission might be a way for the gravid mosquito to prime the environment, *i.e.* the breeding water, with her own beneficial bacteria. Indeed, our data seem to indicate that diagonal transmission applies for the mosquito microbiome, although further experimental studies are required to support these observations. Recently, Nichols *et al.* showed that individual females transmit a subset of their microbiota to their offspring, that maternally derived bacteria support larval development, and that adult offspring microbiota remain similar to those of their mothers despite extensive reshaping during larval stages [55]. They referred to it as “vertical transmission” because they used isofemales within their experimental setup. However, in a natural environment, their findings would support the diagonal transmission hypothesis.

We found that the water of the breeding sites and the *Ae. aegypti* microbiome were dominated by Proteobacteria, Actinobacteriota, Bacteroidota, and Firmicutes. These phyla include many bacteria generally found in aquatic and soil environments. Other bacteria were present in the samples but represented only a minor fraction of the community (**Figure 3**). Interestingly, the abundance of the Actinobacteriota, Bacteroidota, and Firmicutes decreased in the later developmental stages while the Proteobacteria became completely dominant in the adult samples. This result is in line with previous studies reporting that the microbiome of adult *Ae. aegypti* from other countries is also dominated by Proteobacteria [27, 40, 42, 43]. With this high degree of agreement across these different studies, we hypothesize that the Proteobacteria play an important role in the development of the mosquito. This seems to be supported by the fact that the 15 most abundant ASVs of the newly emerged adults analysed in our study all belonged to the Proteobacteria (**Figure 4C**), reflecting a consistent profile at adulthood. Moreover, also the absolute abundances of the top 15 ASVs were quite similar across all the adults (**Figure 5A**).

Proteobacteria is a major phylum of gram-negative bacteria. In our study, we identified Gammaproteobacteria as the most abundant class in adult mosquitoes within the Proteobacteria. Although further distinction into different bacterial species is not possible with our 16S V3-V4 dataset, the retention of Proteobacteria as 15 most abundant ASVs in the newly emerged adults across all the different conditions can indicate a selection of these bacteria in the mosquito. On one hand, these bacteria could be more adapted to grow in the ‘hostile’ mosquito gut environment and are therefore more likely to survive after passive selection from the water during larval feeding. In *Anopheles stephensi* mosquitoes, the Malpighian tubules, excretory and osmoregulatory organs that stay intact during metamorphosis, function as a hide-out for *Pseudomonas* bacteria from the continuous regeneration of the gut epithelial lining during mosquito development [56]. A similar mechanism could possibly explain the trans-stadial selection of these 15 most abundant ASVs and other Proteobacteria in our study. On the other hand, Proteobacteria present in the breeding water could be emitting chemical cues which are picked up by *Aedes* females to deposit their eggs at that site. In both cases, the acquired Proteobacteria can have an impact on the development and/or later life adult life stages of the mosquito. For instance, Proteobacteria could provide the mosquito with a developmental advantage by providing essential nutrients for the developing mosquito. This is further exemplified by restoring developmental issues of germ-free *Ae. aegypti* after recolonization with *E. coli*, a Gammaproteobacteria member [57]. Also in other mosquito genera, *i.e. Anopheles*, the Gammaproteobacteria (*i.e. Pseudomonas* and *Enterobacter*) were identified as dominant and important members of the mosquito microbiome [35, 56]. Furthermore, *Serratia* and *Enterobacter* were demonstrated to aid in blood digestion via hemolysin production in *Aedes* mosquitoes [31]. These examples support the hypothesis that female mosquitoes deposit their beneficial Gammaproteobacteria in the breeding water to facilitate the immature life stages of the offspring mosquito by providing them with developmental benefits.

Although the samples in all examined public studies were also dominated by Proteobacteria (see **Supplemental Figures 6-9**), we did not observe a convergence towards a small set of ASVs in the adult samples of these public datasets (ASVs were independently identified between studies). This can be explained by the sampling methods used in the analysed datasets. These were not focused on sampling each life stage from exactly the same habitat (*i.e.* same breeding container on the same date) as was done in our study. In addition, studies with more than two life stages had rather limited samples [27, 43] and the Hery study [42] did not contain any adult samples, while also not having any water and larvae samples from the same source. This highlights the unique advantage of our sampling procedure (**Figure 1**) and the necessity to confirm our results in future studies with similar sampling methods that can track the abundance of ASVs across the maturation of a mosquito.

On the other hand, limitations of our study are the low sample size per condition (three replicates) and the limited sampling locations (two locations) which may not be representative of the totality of the microbiome of *Ae. aegypti* from Burkina Faso. Additional sampling over a wider geographical distribution and over multiple timepoints (more seasons, more years) might allow the evaluation of the consistency of the microbiome in *Aedes* mosquitoes. Furthermore, we did not include mosquito eggs and only included newly emerged adults which does not allow to see what the effect of nectar and blood feeding is on the composition of the microbiome in adult mosquitoes. Finally, we performed 16S sequencing on the V3-V4 regions which limits the bacterial identification to genus level at best and thus could not give us more information on which bacterial species or strains are present.

## Conclusion

This study profiled the bacterial communities over different mosquito life stages in field *Ae. aegypti* females from Burkina Faso. We demonstrated a decrease in both the bacterial diversity and the bacterial load over the different developmental stages, with the highest diversity and load observed in the breeding water and larvae samples whereas adult mosquitoes possessed the lowest diversity and load. Interestingly, a strong selection of Proteobacteria (more specifically Gammaproteobacteria) in the adult females was identified. The presence of Proteobacteria was demonstrated to be consistent over all adult samples and was irrespective of environmental variables including breeding containers (tire or plastic container) and geographical sampling site (urban or peri-urban site), suggesting they may be transmitted diagonally.

## Acknowledgements

We thank Honoré Badolo for his assistance for the mosquito sampling in the different sites. We also thank the population from the sampling sites who allowed us to collect mosquito samples.

The geographic shapefile for Figure 1A was provided by the Institut Géographique du Burkina (https://data.humdata.org/dataset/cod-ab-bfa) and mosquito vector illustrations for Figure 1B and Supplemental Figure 10 were used from NIAID NIH BIOART Source.

## Author contributions

A.S, L.W, A.B and L.D conceived and organized the study. A.S conducted the experiments, performed basic analysis and wrote the manuscript. L.D.C generated the ASVs table, cleaned the datasets, performed statistical analyses, generated the plots and wrote the manuscript. K.T wrote the manuscript. J.M and L.D reviewed the manuscript. All authors read and approved the manuscript.

## Data availability

Raw sequence data is accessible on SRA under BioProject PRJNA1301862. Scripts used to analyse the data are available at <u>github.com/absanon/2024_XXX_MICROBIOME_LIFE_STAGE</u>.

## Consent for publication

Not applicable.

## Ethics approval and consent to participate

Not applicable.

## Supplementary files

**Supplemental Figure 1:**
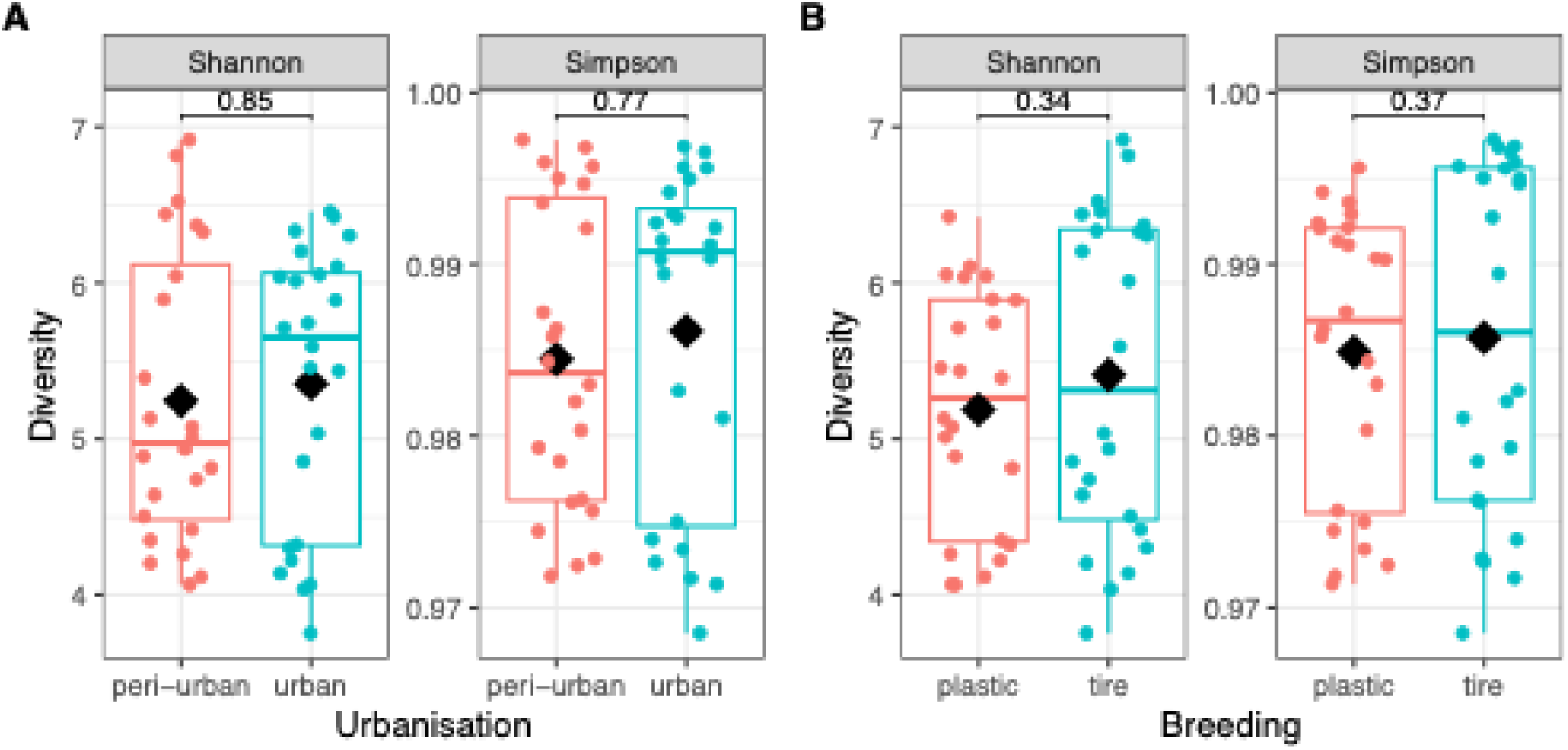
Shannon and Simpson diversity of samples across the two locations (A) and different types of breeding materials (B). The boxplots show the median, 0.25 and 0.75 quantiles, while means for each group are shown by black diamonds. Differences between mean alpha diversity were tested with Wilcoxon tests. P-values are shown between the tested groups.

**Supplemental Figure 2:**
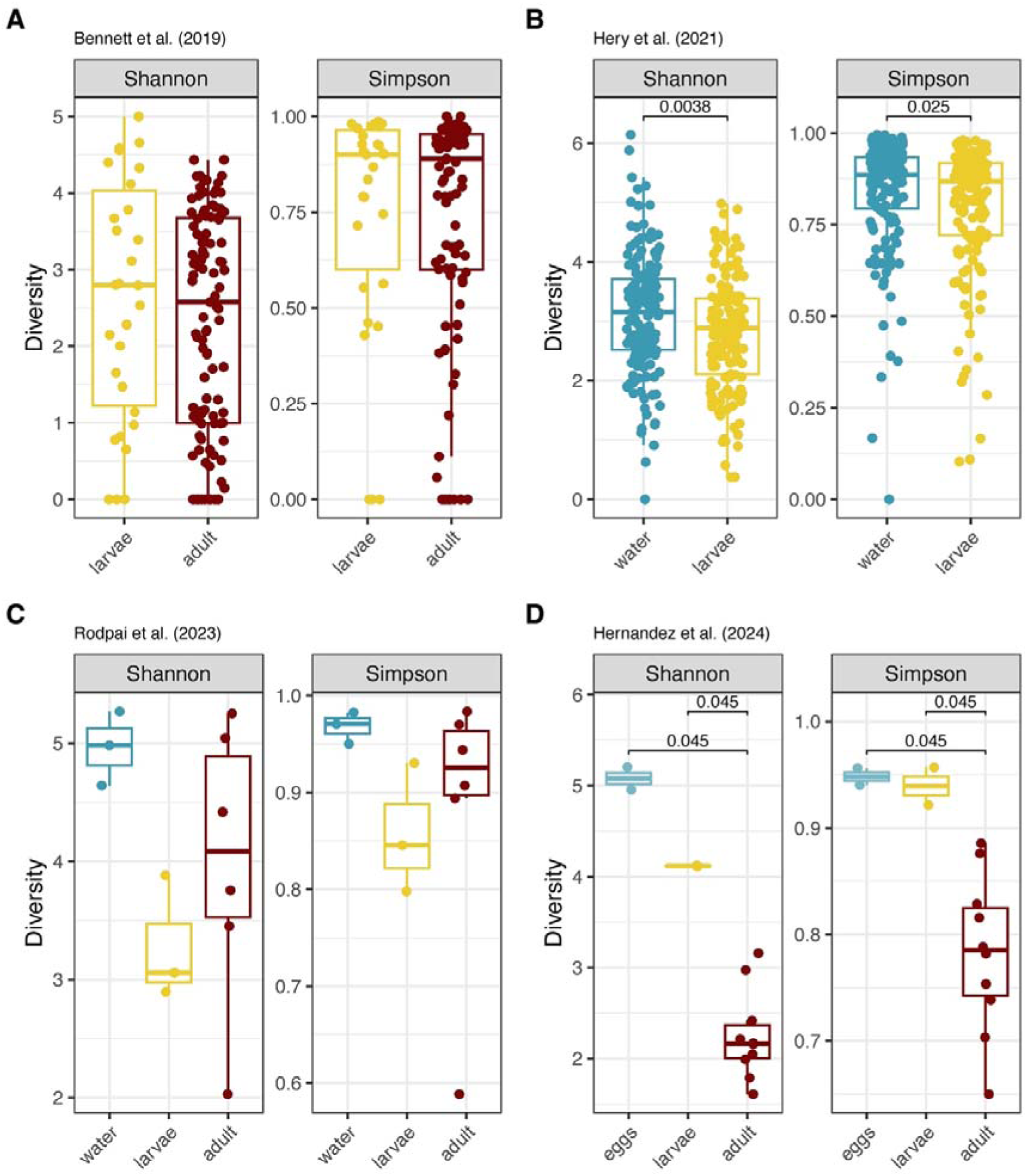
Shannon and Simpson diversity across the different life stages in each study. Differences between mean alpha diversity were tested with Wilcoxon tests and a Benjamini-Hochberg FDR for multiple testing. Only significant differences between groups are shown on top of the boxplots.

**Supplemental Figure 3:**
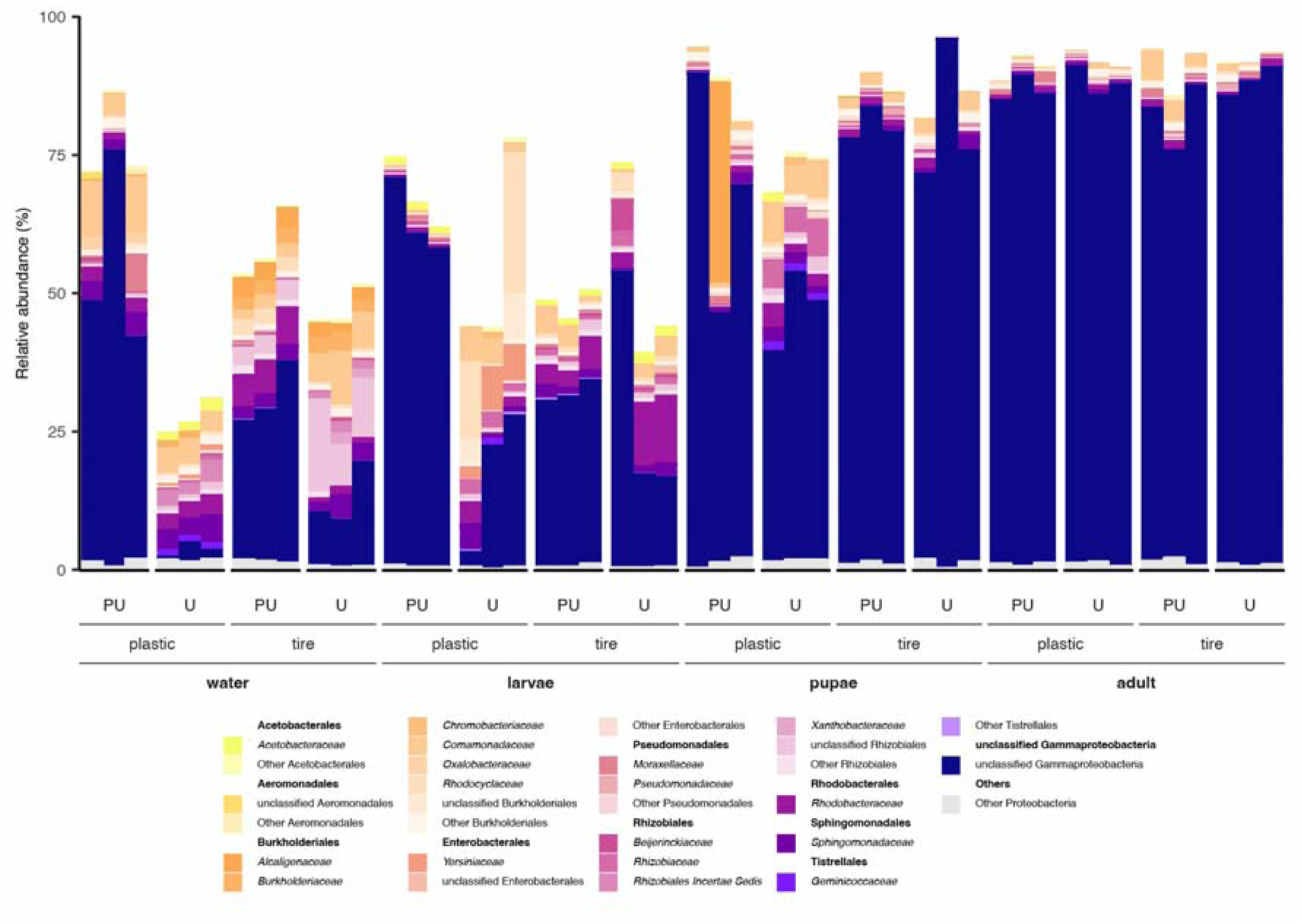
Relative abundance of the *Proteobacteria* phylum per breeding material, sampling sites, and developmental stage. Families have different shades of the same colour if they belong to the same order.

**Supplemental Figure 4:**
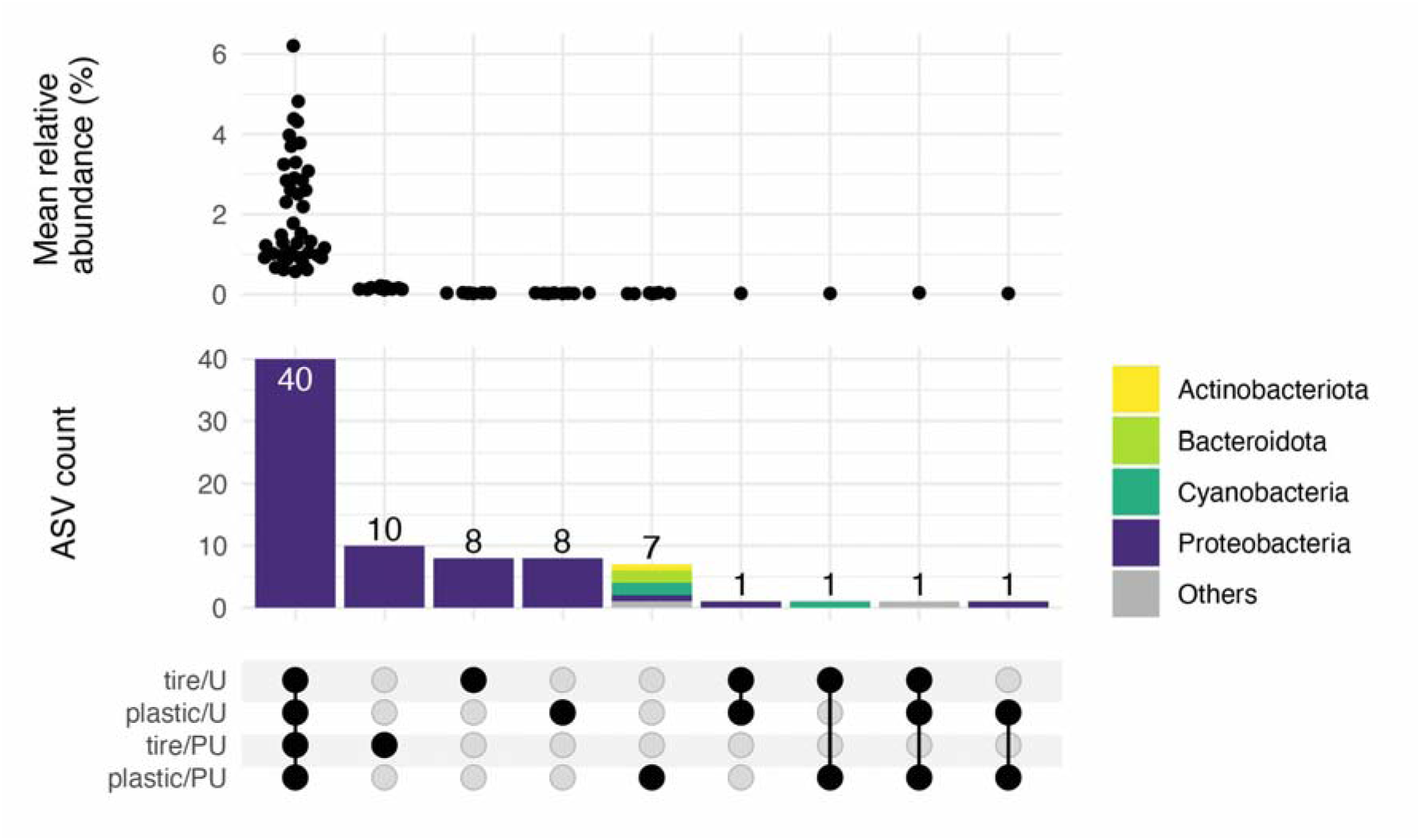
Upset plot showing the distribution of the 50 most abundant ASVs in the adults from each sampling combination (plastic material in urban area, tire in peri-urban area, etc.). On top of the plot the mean relative abundance of each ASV across all considered sampling combinations is given.

**Supplemental Figure 5:**
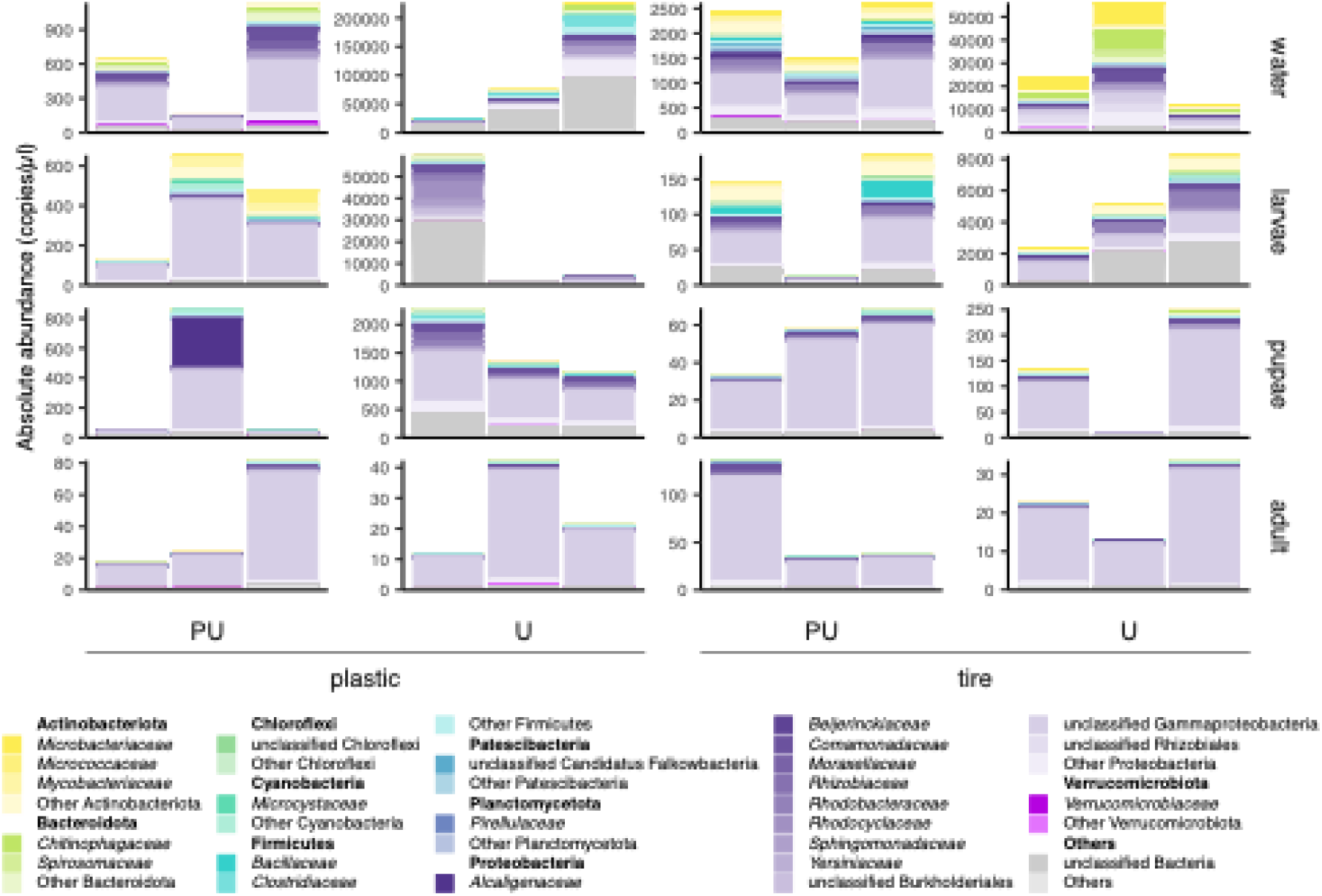
Absolute abundances of the bacterial families based on 16S qPCR. The y-axis shows the number of 16S copies/µl and is scaled to the highest number of copies for each developmental stage with each combination of breeding material and location.

**Supplemental Figure 6:**
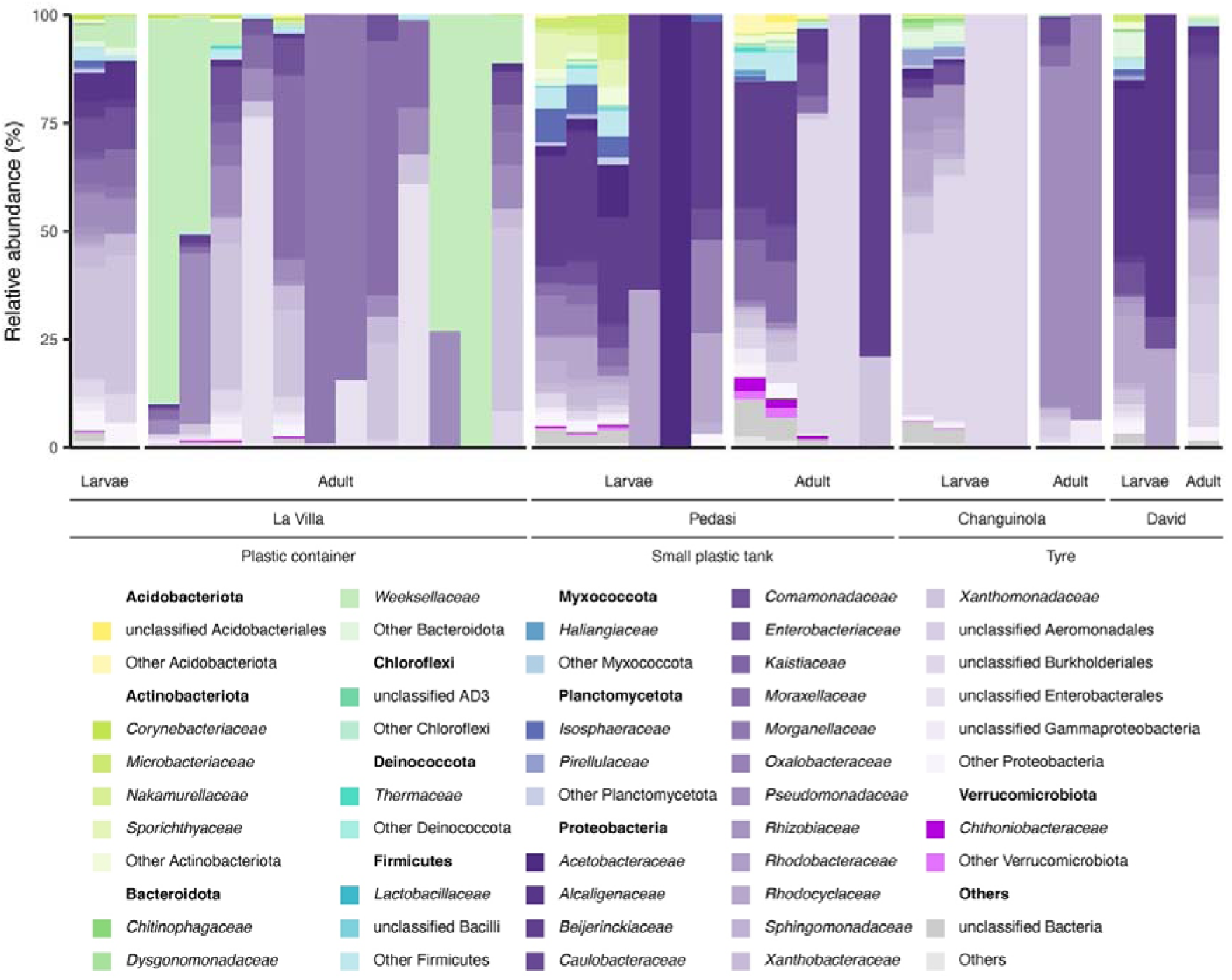
Relative abundance of bacterial families in mosquito samples from Bennett *et al.* (2019).

**Supplemental Figure 7:**
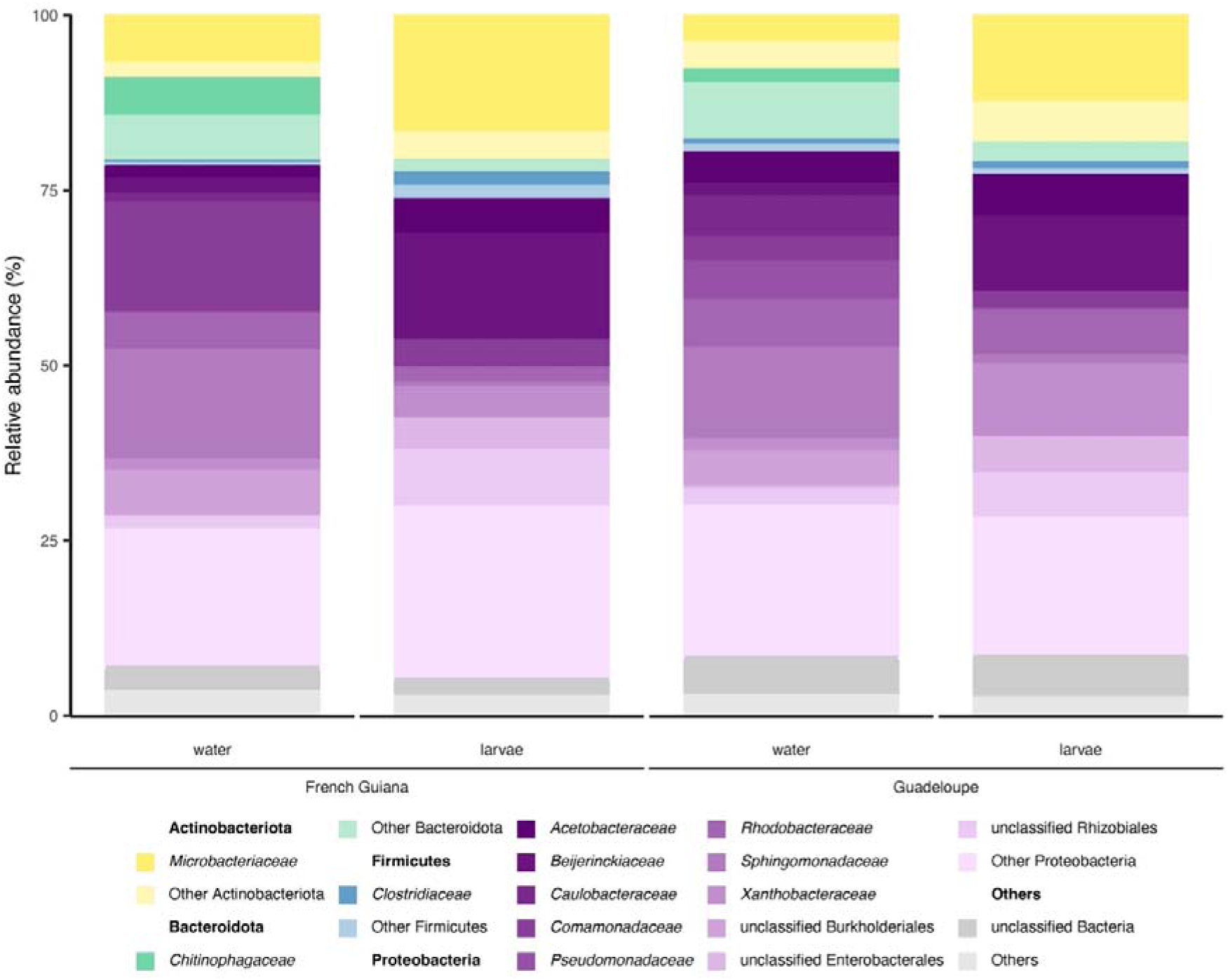
Relative abundance of bacterial families in mosquito samples from Hery *et al.* (2021).

**Supplemental Figure 8:**
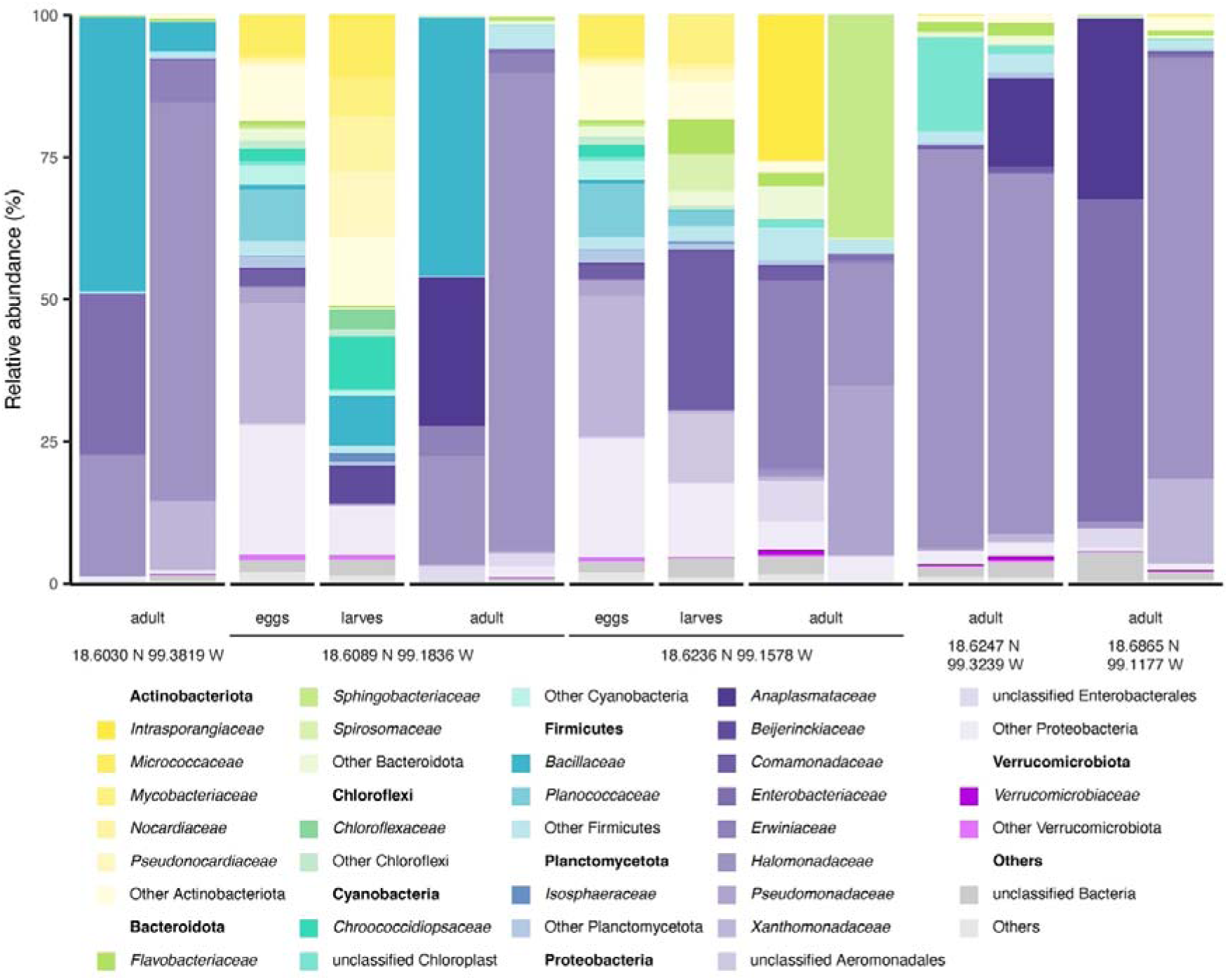
Relative abundance of bacterial familes in mosquito samples from Hernandez *et al.* (2024).

**Supplemental Figure 9:**
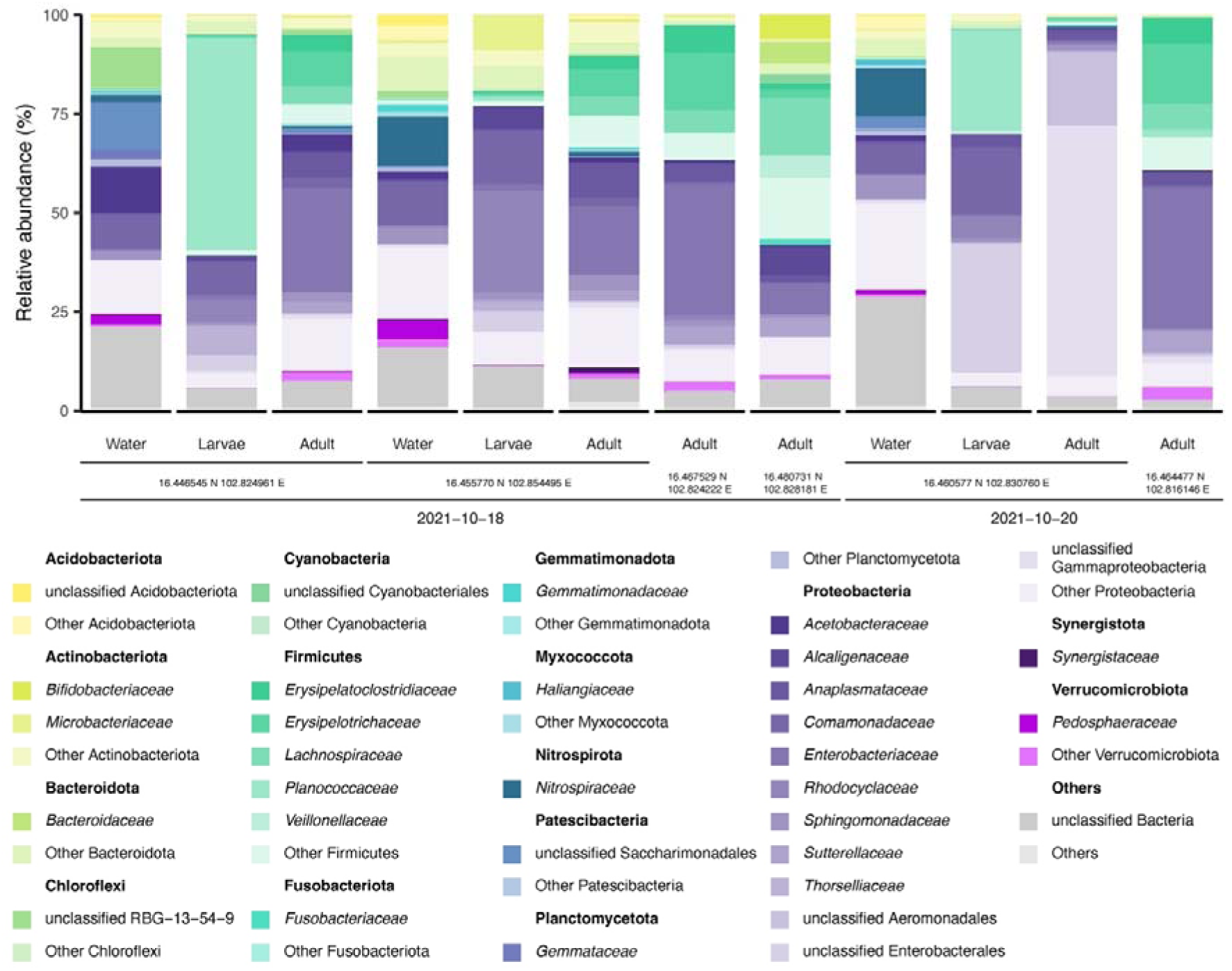
Relative abundance of bacterial families in mosquito samples from Rodpai *et al.* (2023).

**Supplemental Figure 10:**
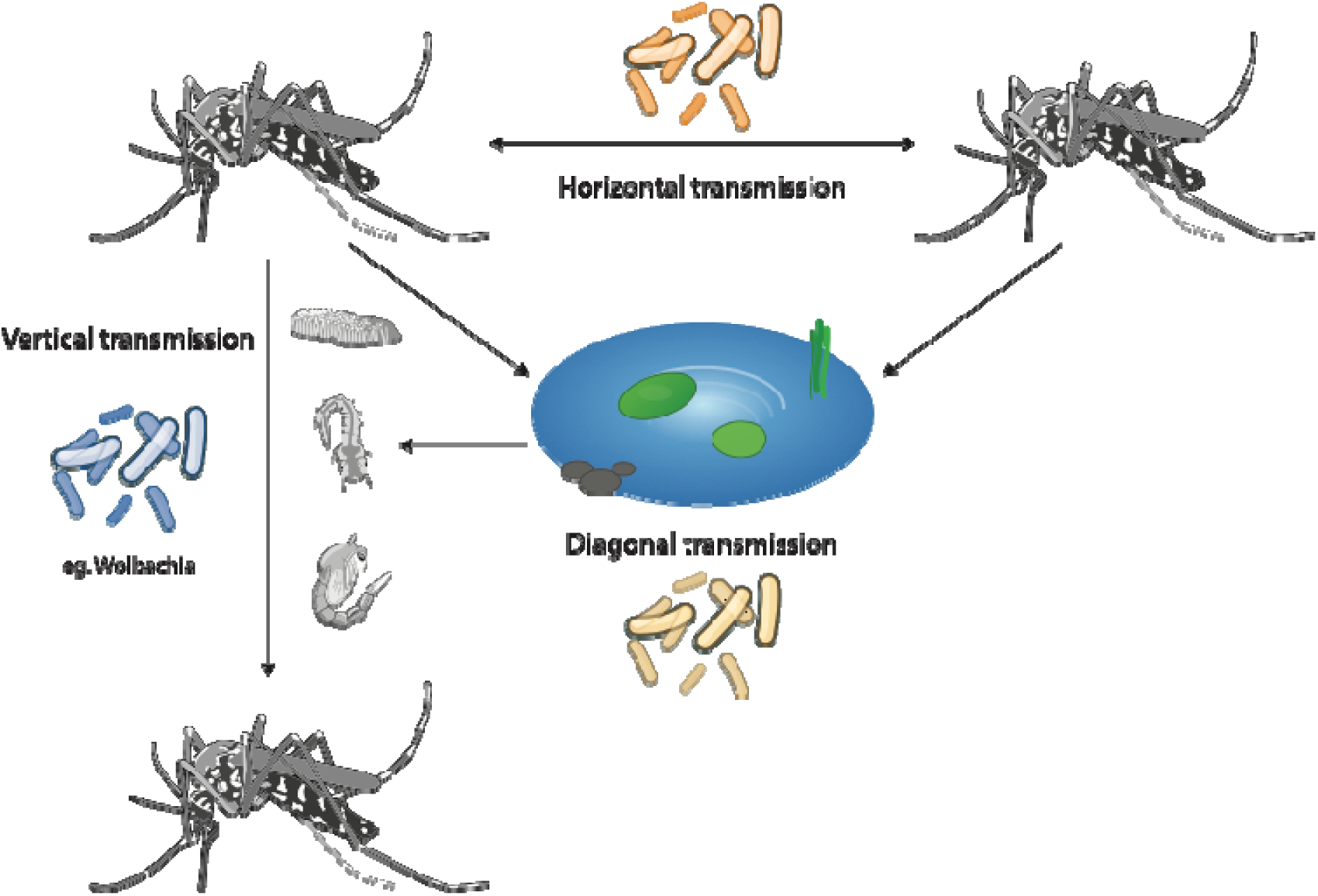
Proposed transmission routes shaping microbiome assembly in mosquitoes. Vertical transmission confers direct transmission from parent to offspring (e.g. Wolbachia is transmitted maternally by infecting germline stem cells in the ovary). Horizontal transmission occurs through environmental exposure, mating, or feeding. Diagonal transmission, on the other hand, constitutes the priming of the breeding environment by female mosquitoes during oviposition, thereby indirectly transferring bacteria to both their offspring and larvae from other females

## References

1. Souza-Neto JA, Powell JR, Bonizzoni M. Aedes aegypti vector competence studies: A review. Infect Genet Evol J Mol Epidemiol Evol Genet Infect Dis. 2019;67:191–209. 10.1016/J.MEEGID.2018.11.009.

2. Badolo A, Sombié A, Yaméogo F, Wangrawa DW, Sanon A, Pignatelli PM, et al. First comprehensive analysis of Aedes aegypti bionomics during an arbovirus outbreak in west Africa: Dengue in Ouagadougou, Burkina Faso, 2016–2017. PLoS Negl Trop Dis. 2022;16:e0010059–e0010059. 10.1371/JOURNAL.PNTD.0010059.

3. Rose NH, Sylla M, Badolo A, Lutomiah J, Ayala D, Aribodor OB, et al. Climate and Urbanization Drive Mosquito Preference for Humans. Curr Biol CB. 2020;30:3570–3579.e6. 10.1016/J.CUB.2020.06.092.

4. Shaw WR, Catteruccia F. Vector biology meets disease control: using basic research to fight vector-borne diseases. Nat Microbiol. 2019;4:20–34. 10.1038/S41564-018-0214-7.

5. Badolo A, Sombié A, Pignatelli PM, Sanon A, Yaméogo F, Wangrawa DW, et al. Insecticide resistance levels and mechanisms in Aedes aegypti populations in and around Ouagadougou, Burkina Faso. PLoS Negl Trop Dis. 2019;13:e0007439–e0007439. 10.1371/JOURNAL.PNTD.0007439.

6. Accoti A, Quek S, Vulcan J, Cansado-Utrilla C, Anderson ER, Abu AEI, et al. Variable microbiomes between mosquito lines are maintained across different environments. PLoS Negl Trop Dis. 2023;17. 10.1371/JOURNAL.PNTD.0011306.

7. Bando H, Okado K, Guelbeogo WM, Badolo A, Aonuma H, Nelson B, et al. Intra-specific diversity of Serratia marcescens in Anopheles mosquito midgut defines Plasmodium transmission capacity. Sci Rep. 2013;3. 10.1038/SREP01641.

8. Clements AN. The biology of mosquitoes. Volume 1: development, nutrition and reproduction. London: Chapman & Hall; 1992. 10.1079/9780851993744.0000.

9. Dong Y, Manfredini F, Dimopoulos G. Implication of the Mosquito Midgut Microbiota in the Defense against Malaria Parasites. PLOS Pathog. 2009;5:e1000423–e1000423. 10.1371/JOURNAL.PPAT.1000423.

10. Merritt RW, Dadd RH, Walker ED. Feeding behavior, natural food, and nutritional relationships of larval mosquitoes. Annu Rev Entomol. 1992;37:349–74. 10.1146/ANNUREV.EN.37.010192.002025.

11. Ricci I, Valzano M, Ulissi U, Epis S, Cappelli A, Favia G. Symbiotic control of mosquito borne disease. Pathog Glob Health. 2012;106:380–5. 10.1179/2047773212Y.0000000051.

12. Hyde J, Correa MA, Hughes GL, Steven B, Brackney DE. Limited influence of the microbiome on the transcriptional profile of female Aedes aegypti mosquitoes. Sci Rep 2020 101. 2020;10:1–12. 10.1038/s41598-020-67811-y.

13. Hegde S, Rasgon JL, Hughes GL. The microbiome modulates arbovirus transmission in mosquitoes. Curr Opin Virol. 2015;15:97–102. 10.1016/j.coviro.2015.08.011.

14. Scolari F, Casiraghi M, Bonizzoni M. Aedes spp. and Their Microbiota: A Review. Front Microbiol. 2019;10. 10.3389/fmicb.2019.02036.

15. Wang J, Gao L, Aksoy S. Microbiota in disease-transmitting vectors. Nat Rev Microbiol 2023 219. 2023;21:604–18. 10.1038/s41579-023-00901-6.

16. Gao H, Cui C, Wang L, Jacobs-Lorena M, Wang S. Mosquito Microbiota and Implications for Disease Control. Trends Parasitol. 2020;36:98–111. 10.1016/J.PT.2019.12.001.

17. Zhang L, Wang D, Shi P, Li J, Niu J, Chen J, et al. A naturally isolated symbiotic bacterium suppresses flavivirus transmission by Aedes mosquitoes. Science. 2024;384:eadn9524. 10.1126/science.adn9524.

18. Girard M, Martin E, Vallon L, Raquin V, Bellet C, Rozier Y, et al. Microorganisms Associated with Mosquito Oviposition Sites: Implications for Habitat Selection and Insect Life Histories. Microorganisms. 2021;9:1589. 10.3390/microorganisms9081589.

19. Day JF. Mosquito Oviposition Behavior and Vector Control. Insects. 2016;7:65. 10.3390/insects7040065.

20. Boissière A, Tchioffo MT, Bachar D, Abate L, Marie A, Nsango SE, et al. Midgut Microbiota of the Malaria Mosquito Vector Anopheles gambiae and Interactions with Plasmodium falciparum Infection. PLOS Pathog. 2012;8:e1002742. 10.1371/journal.ppat.1002742.

21. Steven B, Hyde J, LaReau JC, Brackney DE. The Axenic and Gnotobiotic Mosquito: Emerging Models for Microbiome Host Interactions. Front Microbiol. 2021;12. 10.3389/fmicb.2021.714222.

22. Coon KL, Brown MR, Strand MR. Mosquitoes host communities of bacteria that are essential for development but vary greatly between local habitats. Mol Ecol. 2016;25:5806–26. 10.1111/MEC.13877.

23. Dada N, Jumas-Bilak E, Manguin S, Seidu R, Stenström TA, Overgaard HJ. Comparative assessment of the bacterial communities associated with Aedes aegypti larvae and water from domestic water storage containers. Parasit Vectors. 2014;7:1– 12. 10.1186/1756-3305-7-391/FIGURES/6.

24. Juma EO, Allan BF, Kim C-H, Stone C, Dunlap C, Muturi EJ. The larval environment strongly influences the bacterial communities of Aedes triseriatus and Aedes japonicus (Diptera: Culicidae). Sci Rep. 2021;11:7910. 10.1038/s41598-021-87017-0.

25. Scolari F, Sandionigi A, Carlassara M, Bruno A, Casiraghi M, Bonizzoni M. Exploring Changes in the Microbiota of Aedes albopictus: Comparison Among Breeding Site Water, Larvae, and Adults. Front Microbiol. 2021;12:624170–624170. 10.3389/FMICB.2021.624170/BIBTEX.

26. Capinera JL. Encyclopedia of Entomology. Springer Science & Business Media; 2008.

27. Hernández AM, Alcaraz LD, Hernández-Álvarez C, Romero MF, Jara-Servín A, Barajas H, et al. Revealing the microbiome diversity and biocontrol potential of field Aedes ssp.: Implications for disease vector management. PLOS ONE. 2024;19:e0302328–e0302328. 10.1371/JOURNAL.PONE.0302328.

28. Villegas LEM, Radl J, Dimopoulos G, Short SM. Bacterial communities of Aedes aegypti mosquitoes differ between crop and midgut tissues. PLoS Negl Trop Dis. 2023;17:e0011218. 10.1371/journal.pntd.0011218.

29. Viafara-Campo JD, Vivero-Gómez RJ, Fernando-Largo D, Manjarrés LM, Moreno-Herrera CX, Cadavid-Restrepo G. Diversity of Gut Bacteria of Field-Collected Aedes aegypti Larvae and Females, Resistant to Temephos and Deltamethrin. Insects. 2025;16:181. 10.3390/insects16020181.

30. Dickson LB, Jiolle D, Minard G, Moltini-Conclois I, Volant S, Ghozlane A, et al. Carryover effects of larval exposure to different environmental bacteria drive adult trait variation in a mosquito vector. Sci Adv. 2017;3:e1700585. 10.1126/sciadv.1700585.

31. Gaio A de O, Gusmão DS, Santos AV, Berbert-Molina MA, Pimenta PF, Lemos FJ. Contribution of midgut bacteria to blood digestion and egg production in Aedes aegypti (diptera: culicidae) (L.). Parasit Vectors. 2011;4:105. 10.1186/1756-3305-4-105.

32. Gusmão DS, Santos AV, Marini DC, Bacci M, Berbert-Molina MA, Lemos FJA. Culture-dependent and culture-independent characterization of microorganisms associated with *Aedes aegypti* (Diptera: Culicidae) (L.) and dynamics of bacterial colonization in the midgut. Acta Trop. 2010;115:275–81. 10.1016/j.actatropica.2010.04.011.

33. Guégan M, Tran Van V, Martin E, Minard G, Tran F-H, Fel B, et al. Who is eating fructose within the Aedes albopictus gut microbiota? Environ Microbiol. 2020;22:1193–206. 10.1111/1462-2920.14915.

34. Gimonneau G, Tchioffo MT, Abate L, Boissière A, Awono-Ambéné PH, Nsango SE, et al. Composition of *Anopheles coluzzii* and *Anopheles gambiae* microbiota from larval to adult stages. Infect Genet Evol. 2014;28:715–24. 10.1016/j.meegid.2014.09.029.

35. Galeano-Castañeda Y, Bascuñán P, Serre D, Correa MM. Trans-stadial fate of the gut bacterial microbiota in *Anopheles albimanus*. Acta Trop. 2020;201:105204. 10.1016/j.actatropica.2019.105204.

36. Wang Y, Iii TMG, Kukutla P, Yan G, Xu J. Dynamic Gut Microbiome across Life History of the Malaria Mosquito Anopheles gambiae in Kenya. PLOS ONE. 2011;6:e24767. 10.1371/journal.pone.0024767.

37. Nichols HL, Coon KL. Leveraging microbial ecology for mosquito-borne disease control. Trends Parasitol. 2025. 10.1016/j.pt.2025.06.010.

38. Carvajal-Lago L, Ruiz-López MJ, Figuerola J, Martínez-de la Puente J. Implications of diet on mosquito life history traits and pathogen transmission. Environ Res. 2021;195:110893. 10.1016/j.envres.2021.110893.

39. Wang X, Liu T, Wu Y, Zhong D, Zhou G, Su X, et al. Bacterial microbiota assemblage in Aedes albopictus mosquitoes and its impacts on larval development. Mol Ecol. 2018;27:2972–85. 10.1111/mec.14732.

40. Bennett KL, Gómez-Martínez C, Chin Y, Saltonstall K, McMillan WO, Rovira JR, et al. Dynamics and diversity of bacteria associated with the disease vectors Aedes aegypti and Aedes albopictus. Sci Rep 2019 91. 2019;9:1–12. 10.1038/s41598-019-48414-8.

41. Coon KL, Vogel KJ, Brown MR, Strand MR. Mosquitoes rely on their gut microbiota for development. Mol Ecol. 2014;23:2727–39. 10.1111/mec.12771.

42. Hery L, Guidez A, Durand AA, Delannay C, Normandeau-Guimond J, Reynaud Y, et al. Natural Variation in Physicochemical Profiles and Bacterial Communities Associated with Aedes aegypti Breeding Sites and Larvae on Guadeloupe and French Guiana. Microb Ecol. 2021;81:93–109. 10.1007/S00248-020-01544-3.

43. Rodpai R, Boonroumkaew P, Sadaow L, Sanpool O, Janwan P, Thanchomnang T, et al. Microbiome Composition and Microbial Community Structure in Mosquito Vectors Aedes aegypti and Aedes albopictus in Northeastern Thailand, a Dengue-Endemic Area. Insects. 2023;14. 10.3390/INSECTS14020184.

44. Callahan BJ, McMurdie PJ, Rosen MJ, Han AW, Johnson AJA, Holmes SP. DADA2: High resolution sample inference from Illumina amplicon data. Nat Methods. 2016;13:581–581. 10.1038/NMETH.3869.

45. Murali A, Bhargava A, Wright ES. IDTAXA: A novel approach for accurate taxonomic classification of microbiome sequences. Microbiome. 2018;6:1–14. 10.1186/S40168-018-0521-5/FIGURES/6.

46. Davis NM, Proctor DiM, Holmes SP, Relman DA, Callahan BJ. Simple statistical identification and removal of contaminant sequences in marker-gene and metagenomics data. Microbiome. 2018;6:1–14. 10.1186/S40168-018-0605-2/FIGURES/6.

47. Oksanen J, Simpson GL, Blanchet FG, Kindt R, Legendre P, Minchin PR, et al. vegan: Community Ecology Package. 2024.

48. Schloss PD. Rarefaction is currently the best approach to control for uneven sequencing effort in amplicon sequence analyses. mSphere. 2024;9. 10.1128/MSPHERE.00354-23.

49. Paradis E, Schliep K. ape 5.0: an environment for modern phylogenetics and evolutionary analyses in R. Bioinformatics. 2019;35:526–8. 10.1093/BIOINFORMATICS/BTY633.

50. Nadkarni MA, Martin FE, Jacques NA, Hunter N. Determination of bacterial load by real-time PCR using a broad-range (universal) probe and primers set. Microbiology. 2002;148:257–66. 10.1099/00221287-148-1-257.

51. Moll RM, Romoser WS, Modrakowski MC, Moncayo AC, Lerdthusnee K. Meconial Peritrophic Membranes and the Fate of Midgut Bacteria During Mosquito (Diptera: Culicidae) Metamorphosis. J Med Entomol. 2001;38:29–32. 10.1603/0022-2585-38.1.29.

52. Zouache K, Martin E, Rahola N, Gangue MF, Minard G, Dubost A, et al. Larval habitat determines the bacterial and fungal microbiota of the mosquito vector Aedes aegypti. FEMS Microbiol Ecol. 2022;98. 10.1093/FEMSEC/FIAC016.

53. Mosquera KD, Martínez Villegas LE, Rocha Fernandes G, Rocha David M, Maciel-de-Freitas R, A. Moreira L, et al. Egg-laying by female Aedes aegypti shapes the bacterial communities of breeding sites. BMC Biol. 2023;21:1–15. 10.1186/S12915-023-01605-2/FIGURES/5.

54. Hamel R, Narpon Q, Serrato-Pomar I, Gauliard C, Berthomieu A, Wichit S, et al. West Nile virus is transmitted within mosquito populations through infectious mosquito excreta. bioRxiv. 2024;:2024.01.29.577888-2024.01.29.577888. 10.1101/2024.01.29.577888.

55. Nichols HL, Brettell LE, Saldaña MA, Hegde S, Heinz E, Vaccaro K, et al. Vertical transmission of mosquito microbiota and its effects on offspring development. 2025;:2025.12.16.694692. 10.64898/2025.12.16.694692.

56. Chavshin AR, Oshaghi MA, Vatandoost H, Yakhchali B, Zarenejad F, Terenius O. Malpighian tubules are important determinants of Pseudomonas transstadial transmission and longtime persistence in Anopheles stephensi. Parasit Vectors. 2015;8. 10.1186/s13071-015-0635-6.

57. Correa MA, Matusovsky B, Brackney DE, Steven B. Generation of axenic Aedes aegypti demonstrate live bacteria are not required for mosquito development. Nat Commun. 2018;9:4464. 10.1038/s41467-018-07014-2.

